# Imagery of movements immediately following performance allows learning of motor skills that interfere

**DOI:** 10.1101/299594

**Authors:** Hannah R. Sheahan, James N. Ingram, Goda M. Žalalytė, Daniel M. Wolpert

## Abstract

Motor imagery, that is the mental rehearsal of a motor skill, can lead to improvements when performing the same skill. Here we show a powerful and complementary role, in which motor imagery of movements after actually performing a skill allows learning that is not possible without imagery. We leverage a well-studied motor learning task in which subjects reach in the presence of a dynamic (force-field) perturbation. When two opposing perturbations are presented alternately for the same physical movement, there is substantial interference, preventing any learning. However, when the same physical movement is associated with follow-through movements that differ for each perturbation, both skills can be learned. Here we show that when subjects perform the skill and only imagine the follow-through, substantial learning occurs. In contrast, without such motor imagery there was no learning. Therefore, motor imagery can have a profound effect on skill acquisition even when the imagery is not of the skill itself. Our results suggest that motor imagery may evoke different neural states for the same physical
state, thereby enhancing learning.

## Introduction

The ability to acquire new motor skills without disrupting existing ones is critical to the development of a broad motor repertoire. We have previously suggested that the key to representing multiple motor memories is to have each associated with different neural states, rather than physical states of the body^1^. Specifically, we proposed that when reaching in two opposing force-field environments which alternate randomly from trial to trial, the inability of subjects to learn^2–7^ is due to the fact that each movement is associated with the same neural states. However, contexts which separate neural states for the same physical states should allow learning by enabling the same physical movement to be associated with different motor commands. For example, if each movement through the force-field is part of a larger motor sequence comprised of a different follow-through movement, two opposing perturbations can be learned^1, 6^. As motor preparation is thought to involve setting the initial neural state^8^, just planning different follow-through movements, without execution, results in learning of distinct representations^1^. From this perspective, other behaviours that create different neural states for the same physical states may also enable the learning of distinct motor memories.

Many studies have suggested that imagining a movement and physically executing it may engage similar neural substrates. For example, human neuroimaging research has shown similar motor-related activity when imagining and executing movements^9–12^. Moreover, simply imagining moving a body part increases the EMG response of the associated muscles to TMS over primary motor cortex, suggesting that the circuits involved in action are at least partially active during imagery^13, 14^. Similarly, direct recording of neural populations have recently revealed that when monkeys covertly control a BMI-cursor, the evolution of neural states associated with the preparation and execution of the BMI movements are similar and specific to those observed during the corresponding physical reaches^15^. Given that similar motor cortical dynamics are seen in human and non-human primates^16^, we hypothesized that the same overlap of dynamical neural states may also exist when humans execute or imagine movements. That is, if the neural states of a motor area involved in generating a physical movement can be made different (even partially) by motor imagery, then each of these different neural states could be associated with learning a different motor skill. We hypothesized that imagining movements results in distinct neural states that can drive the formation and retrieval of different motor memories. In contrast to studies of mental rehearsal in which the motor skill is imagined but not performed, here we ask whether performing the skill as part of a larger, imagined motor sequence affects its representation. Specifically, we ask whether two opposing perturbations which would normally interfere, can be learned if each is associated with an imagined follow-through movement. We show that when participants produce the same physical reach, but imagine performing follow-throughs that differ for each field, substantial learning occurs. Moreover, we find that learning under imagery transfers partially to actual movements, suggesting that motor imagery and execution engage overlapping neural states. In contrast, without motor imagery there was no learning. Our results suggest that motor imagery can have a profound effect on skill acquisition and the representation of motor memories, even when the imagery is not of the skill itself.

## Results

Five groups of participants performed a motor learning task. Participants grasped the handle of a robotic interface and made reaching movements from one of four starting locations through a perturbing force field to a central target (see Methods). On exposure trials, the field direction (clockwise or counter-clockwise) was randomly selected on each trial. We associated the direction of the force field with the location of a secondary target which was at ±45° relative to the movement to the central target. The groups differed in whether they were required to continue the reach from the central target to the secondary target and what instructions they were given (Figure 1A).

**Figure 1.**
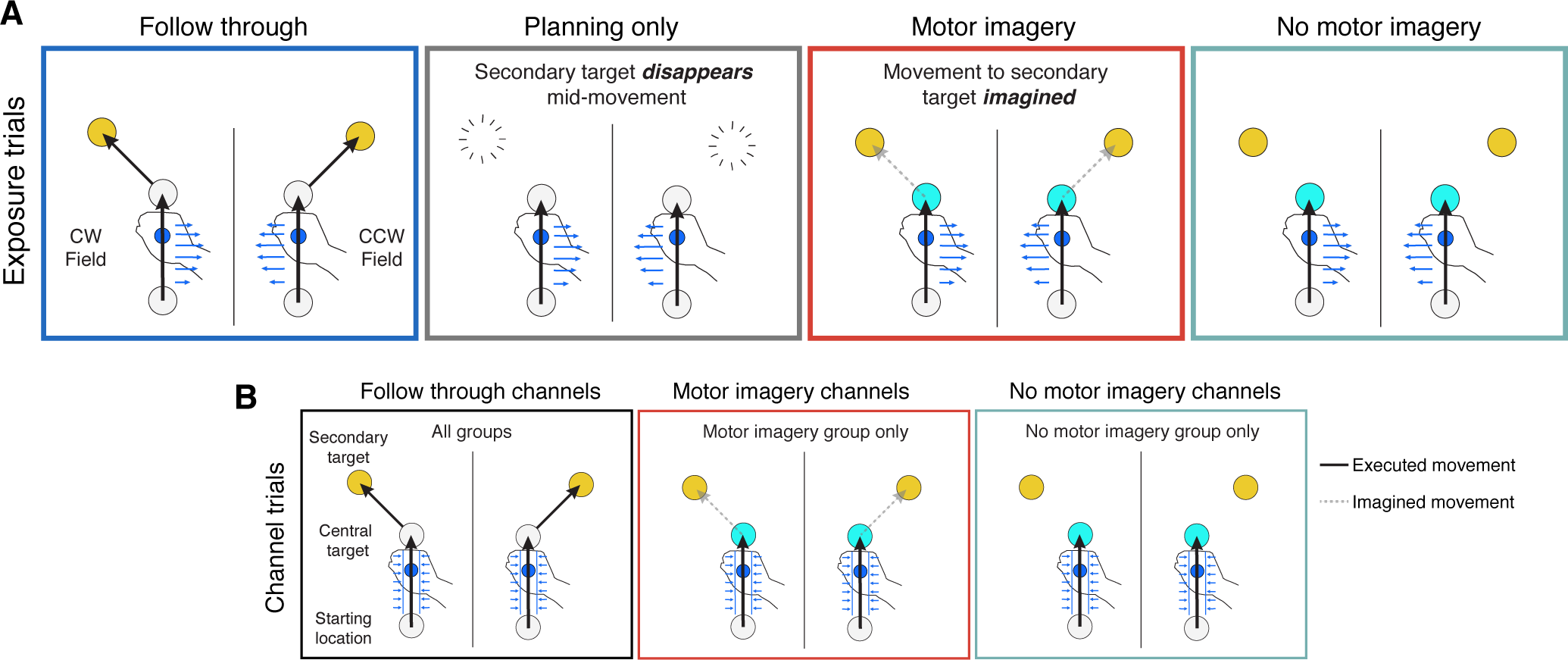
Experimental paradigm. Subjects performed reaching movements that were either (A) exposure trials or (B) channel trials. On all trials a starting location, central target and one secondary target (at either −45° or +45° relative to the initial movement direction) were displayed from the start of the trial. (A) On exposure trials, a velocity-dependent curl force field (blue arrows) was applied on the initial movement. The field direction, clockwise (CW) or counter-clockwise (CCW) was determined by the secondary target location. The exposure trials varied across the groups. The Follow through group continued the initial movement to the secondary target (null field as in channel trials). For the Planning only group, the secondary target disappeared late in the initial movement and they were required to stop at the central target. Both the Motor imagery and No-motor imagery groups were cued by a blue central target, displayed from the start of the trial, that they should stop the movement at the central target. In addition, the motor imagery groups were asked to imagine making a movement to the secondary target and press a button when the imagined movement was complete. (B) On follow through channel trials (left), subjects made a movement to the central target followed immediately by a movement to the secondary target. A channel was applied on the initial movement, allowing an assessment of adaptation measured as the forces applied into the channel wall. A null field was applied on the secondary movement. For half of participants in the motor imagery group, we also included channels for imagined follow though trials (middle) at the end of the exposure phase. Likewise, for half of participants in the no-motor imagery group we included channels for movements just to the central target (right). Note that for clarity in all panels the trials for the two different secondary targets are shown separated, but in the experiment the starting and central targets were in identical locations so that the initial movements were the same. In the experiment there were 4 possible starting locations but for clarity we display only one.

During the exposure phase, we interspersed exposure trials with channel trials, in which the movement was confined to a simulated mechanical channel from the start to central target. For all groups, on these channel trials subjects made a follow through movement to the secondary target which was unconstrained (Figure 1B, left). Note that the simulated channel did not expose subjects to the force field and therefore learning was not possible on these trials. The channel trials allowed us to measure predictive force compensation (the force applied by the participant into the channel wall, expressed as percent adaptation) on the initial movement, independent from factors such as co-contraction^17, 18^. On non-channel trials, we also calculated the maximum perpendicular error (MPE) of the hand path to the central target, which is a measure of the kinematic error of the movement.

On exposure trials, the first group of participants were required to make a second unperturbed follow through movement to the secondary target immediately after arriving at the central target (Figure 1A, Follow through). Importantly, this follow through movement was predictive of the field direction. The second group planned the follow through, but never executed it on exposure trials (Figure 1A, Planning only). That is, the secondary target was displayed from the start of the trial but vanished during the initial movement indicating that the subject should terminate the movement at the central target. To encourage the planning of the entire movement, this group (like all other groups) also made full follow through movements on channel trials (Figure 1B, left).

Both these groups showed significant learning of the two force fields (adaptation increases of 42.9 ± 7.5%, t(7) = 5.92, p = 5.9e-4 and 41.9 ± 4.8%, t(7) = 9.87, p = 2.3e-5 for the follow though and planning groups, respectively), reaching approximately 40% of full compensation (Figure 2A, blue and grey). Figure 2B shows the MPE across the exposure blocks and Figure 3 shows the hand paths for key phases of the experiment for all groups. Moreover, both these groups showed significant aftereffects when the force field was removed during the post-exposure phase (difference in MPE between pre- and post-exposure; 0.94 ± 0.14 cm, t(7) = 7.28, p = 1.7e-4, and 0.78 ± 0.17 cm, t(7) = 5.05, p = 0.0015 for each group respectively). These first two groups included data from six subjects from a previously published study^1^, together with two additional subjects in each group, to provide a baseline for the new groups. The amount of learning in our follow through and planning only groups is similar to that observed in both a previous follow-through study^19^ as well as when learning opposing force fields which are associated with other contextual cues^5^.

**Figure 2.**
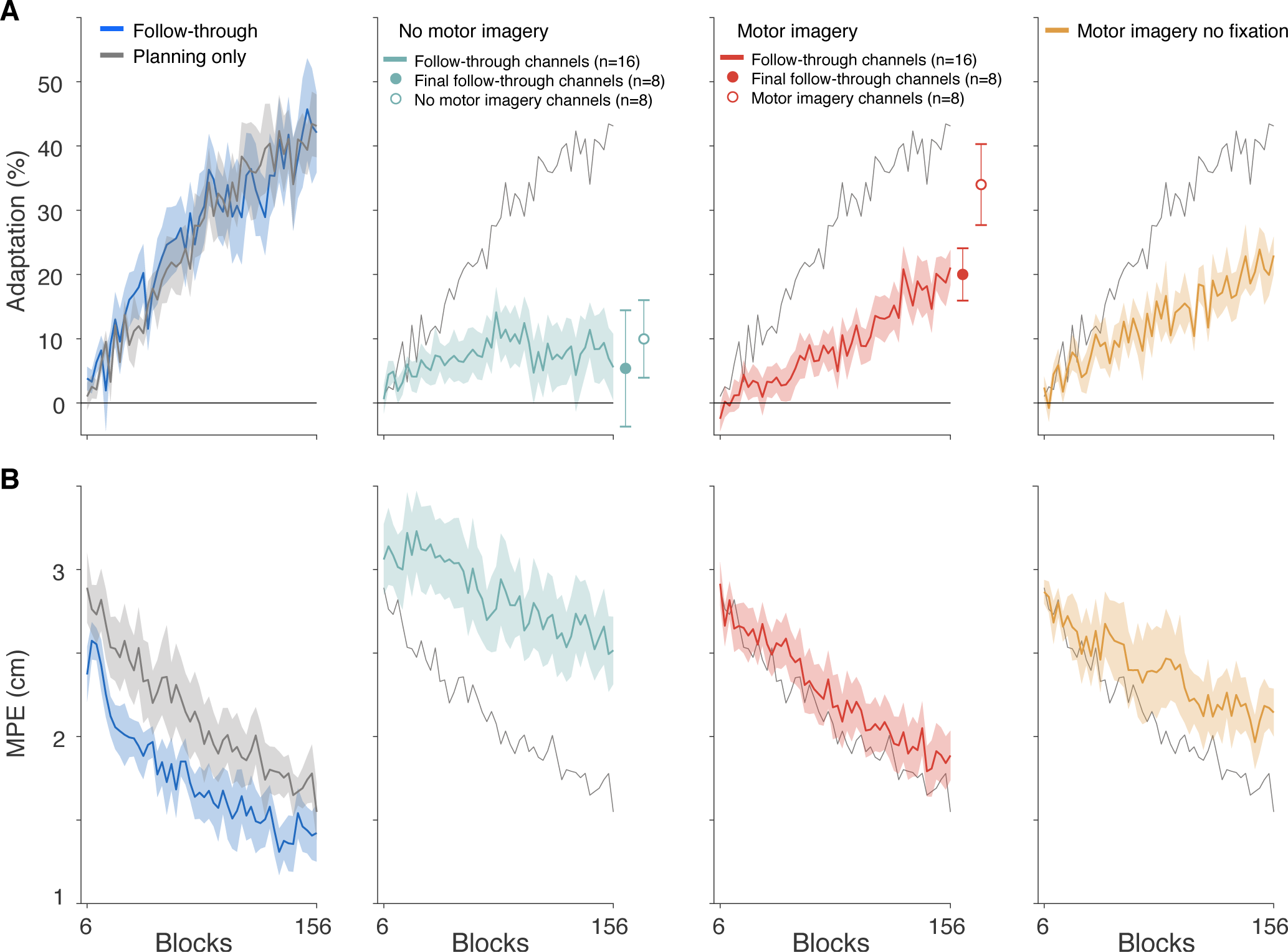
Motor imagery of follow-through leads to adaptation. (A) Adaptation in the exposure phase, measured on channel trials which were full follow through movements in all groups (Figure 1B, left). The solid circles to the right of the imagery and no-imagery learning curves are the final adaptation on full follow-through channel trials of the participants who subsequently performed the probe phase (half the subjects). The unfilled circles show the adaptation measured on motor imagery- or no-motor imagery channel trials (Figure 1B, middle and right) in the same subjects as the solid circles. (B) Maximum perpendicular error (MPE) measured on exposure trials. Data show mean ± s.e. across participants (for display the 150 exposure blocks were averaged over consecutive blocks of 3 to produce 50 bins) in the exposure phase. For comparison, the mean adaptation and MPE for the planning only group (grey) are repeated on all panels.

**Figure 3.**
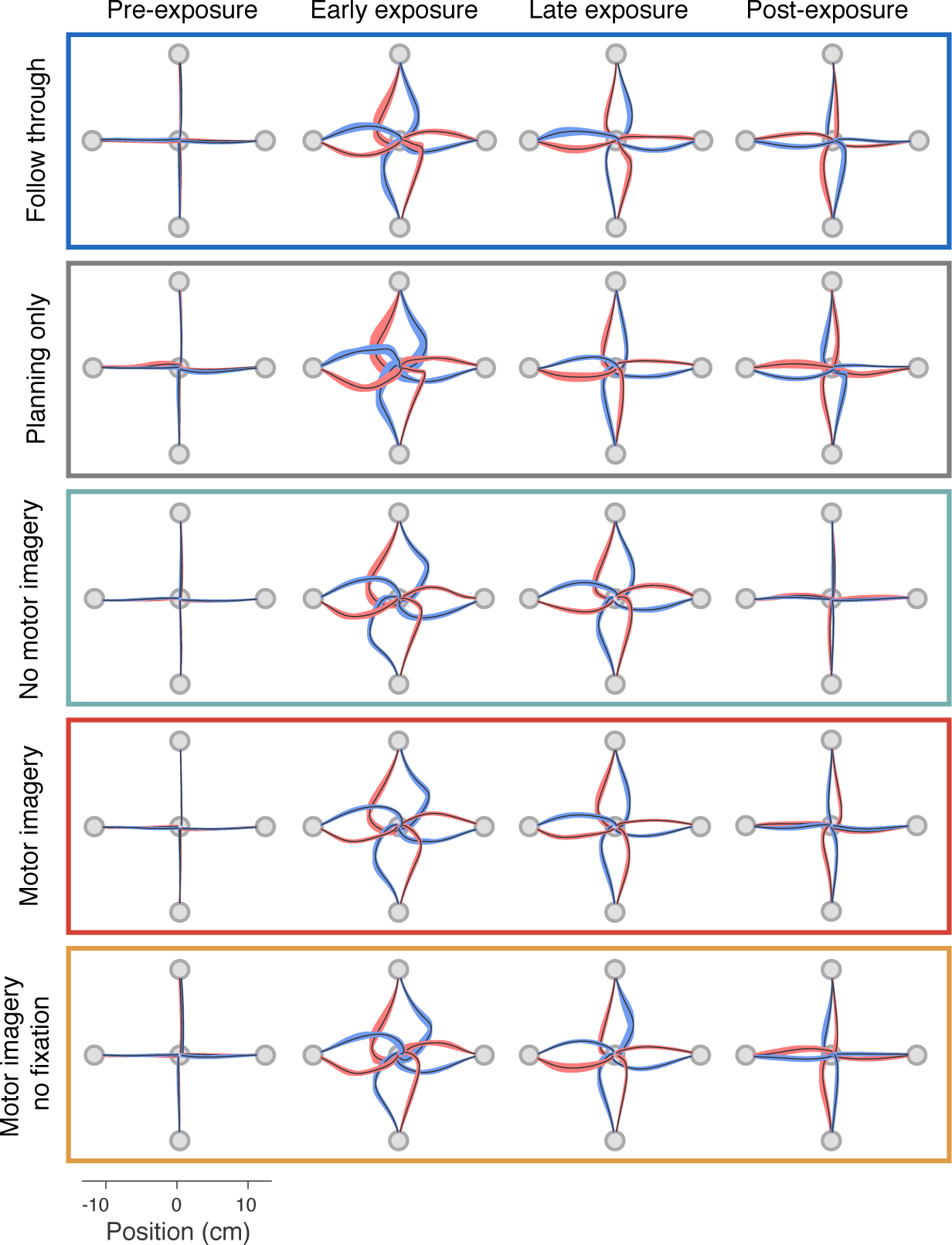
Kinematics across groups for different phases of the experiment. Hand paths are shown from each starting location to the central target for all groups across four different phases of the experiment. Paths show the mean *±* s.e. across participants, for pre-exposure (last block), early exposure (first block), late exposure (last block), and post-exposure (first block). The colors indicate the field direction (blue for CW and red for CCW).

To assess whether motor imagery, like planning, is sufficient to separate motor memories, we compared a no-imagery and an imagery group (Figure 1A). As in the follow through and planning only groups, on channel trials the central target was grey, and participants produced a full follow-through movement. In contrast to the follow through and planning only groups, on exposure trials the central target was blue, such that subjects knew from the start of the trial that they were required to stop at the central target without making a follow through movement. Both groups maintained fixation on the central target throughout each trial. Critically, the motor imagery group was asked to then imagine making the follow-through movement to the secondary target, whereas the no-imagery group was given no such instructions. Therefore, for the motor imagery group, the imagined follow through movement was specific to the force field. To complete a motor imagery trial, these participants pressed a button with their left hand to indicate when the imagined movement reached the secondary target. Importantly, the button-press was the same for both secondary targets, and was therefore not specific to the force-field direction. In the no-imagery group, there was no button press, but we controlled the time spent at the central target by making participants wait for the average amount of time it took them to execute follow-through movements (on channel trials). Consequently, the amount of time spent waiting at the central target did not differ between the imagery and no imagery groups (difference of 72 ± 41 ms, t(30) = 1.72, p = 0.096). After the exposure phase, a subgroup of the participants in each group (n=8) performed a post-exposure phase, identical to the two previous groups, so that we could assess aftereffects. The other participants proceeded to a probe phase of the experiment (see below).

Despite knowing prior to movement initiation that the movement would end at the central target, the motor imagery group increased their adaptation (Figure 2A, red; increase of 21.8 ± 3.1%, t(15) = 7.47, p = 2.0e-6) and also produced significant aftereffects (Figure 3 and Figure 4C, red; 0.68 ± 0.20 cm, t(7) = 4.02, p = 0.0051). In contrast, the no-imagery group showed no significant increase in adaptation (Figure 2A, turquoise; increase of 5.1 ± 5.1%, t(15) = 1.21, p = 0.24) and did not produce significant aftereffects (Figure 3 and Figure 4C, turquoise; no motor imagery group 0.17 ± 0.11 cm, t(7) = 1.83, p = 0.11). This suggests that any reduction in MPE during exposure in the no-imagery group was a consequence of co-contraction or other non-specific strategies. The absence of adaptation in the no imagery group is consistent with many studies which have shown that static visual cues are insufficient to reduce interference to opposing force fields^1, 3–6^. Our results suggest that just imagining the follow through movement allows the separation of motor memories.

**Figure 4.**
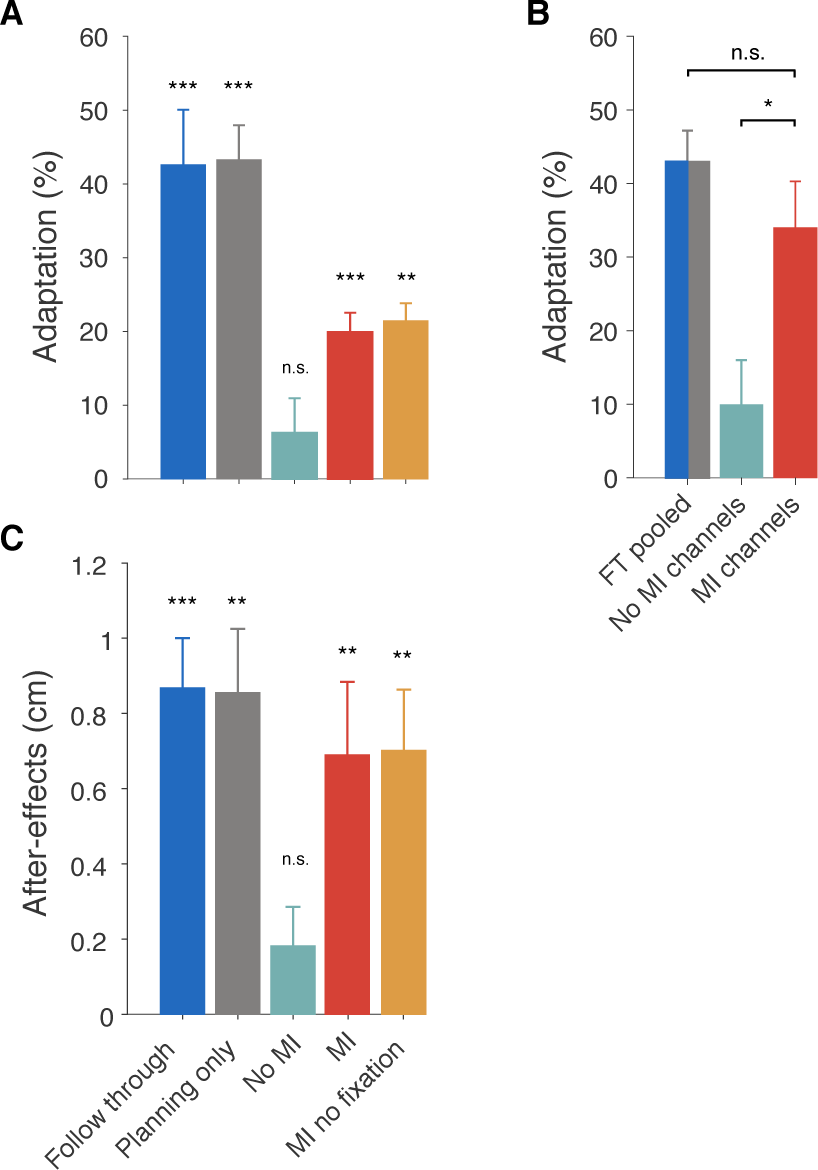
Adaptation and aftereffects. (A) Final adaptation measured on follow through channel trials for all participants (mean ± s.e. of final six blocks of exposure). (B) Comparison of the adaptation between the pooled follow through group and adaptation measured in the subjects who performed the probe phase in the motor imagery and no-imagery groups (first 4 blocks after exposure). (C) MPE during the post-exposure phase (mean ± s.e. of first two blocks) showing aftereffects. Here we consider only the no motor imagery and motor imagery participants who did not perform the probe phase. Therefore all participants shown in (C) experienced the same number of exposure trials before after-effects were assessed. Asterisks show statistical significance of final adaptation level (A) and after-effects (C) compared to pre-exposure, and of differences between groups (B). MI = motor imagery; FT = follow through; n.s. = not significant.

During exposure, adaptation was measured on channel trials with full follow through movements (Figure 1B, left). In the imagery and no-imagery groups, this reflects the transfer of learning from experience of the force field on movements that stop at the central target, to full follow through movements. In order to assess the amount of adaptation on the non-follow through movements themselves (on which the force field was experienced), a subgroup of participants in each of the motor imagery group (subgroup n=8) and no motor imagery group (subgroup n=8) performed an additional phase in which we included channels on non-follow through trials (Figure 1B, middle and right). Final adaptation on follow-through trials was not significantly different between the two subgroups in either the motor imagery group (difference of 0.2%, t(14) = 0.04, p = 0.97, Figure S1) or the no motor imagery group (difference of 1.7%, t(14) = 0.18, p = 0.86, Figure S1). The adaptation measured on motor imagery channel trials (motor imagery group) was 34.0 ± 6.3% (Figure 2A red, motor imagery channels), and no motor imagery channel trials (no motor imagery group) was 10.0 ± 6.0% (Figure 2A, turquoise). There was a significant difference between these groups (Figure 4B;f 24.0 ± 8.7%, t(14) = 2.76, p = 0.015). For comparison, we pooled the follow through and planning only groups (pooled follow through group) based on previous results showing their similar levels of adaptation. We compared the final adaptation in the pooled follow through group with the adaptation measured on motor imagery channel trials in the imagery group (Figure 4B), and found no significant difference (difference in adaptation of 8.9 ± 7.6%, t(22) = 1.20, p = 0.24). This suggests that imagining follow though movements affords similar levels of adaptation as executing or planning to execute them.

In contrast to the motor imagery group, in the full follow through and planning only groups we did not constrain eye movements. To examine whether potential eye movements to the secondary targets could influence learning, we repeated the motor imagery task but without constraining eye movements. This no fixation motor imagery group increased their adaptation (Figure 2A, orange; increase of 17.2 ± 3.1%, t(7) = 4.66, p = 0.0023) and produced significant aftereffects (Figure 3 and Figure 4C, orange; 0.75 ± 0.17 cm, t(7) = 4.98, p = 0.0016). We compared the motor imagery groups with and without fixation, and found no significant effect of fixation on adaptation (difference of 1.5 ± 3.6%, t(22) = 0.375, p = 0.71). This suggests that constraining eye movements did not influence learning.

We compared the final adaptation levels across groups (Figure 4A). The final adaptation in the no imagery group was significantly less (difference of 13.6 ± 5.4%, t(30) = 2.53, p = 0.017) than the motor imagery group, suggesting that imagining follow through movements has a strong effect on learning. In addition, the pooled follow through group had significantly greater adaptation than the motor imagery group (difference of 23.0 ± 5.0%, t(30) = 4.62, p = 6.9e-5) showing that learning under motor imagery does not transfer fully to actual follow throughs. The aftereffects mirror the results seen in the measures of adaptation (Figure 3 and Figure 4C). Shapiro-Wilk tests could not reject the null hypothesis that final adaptation data was normally distributed.

Comparing the chronometrics of imagined and executed movements in the motor imagery groups^20, 21^, there was no significant difference in the durations across participants (Figure 5C difference of 11 ± 3 ms, t(23) = 0.37, p = 0.71), although the variability was higher for the imagined durations. The absolute time difference between executed and imagined follow-throughs in each participant was uncorrelated with their final level of adaptation (r = −0.05, p = 0.802). In addition, there was no correlation between the final level of adaptation in the motor imagery groups and the self-reports from subjects as to how often they remembered to imagine follow-through movements (r = 0.24, p = 0.25), the ease of imagery maintenance (r = 0.14, p = 0.51), or the Motor Imagery Questionnaire (MIQ-RS) scores (r = −0.03, p = 0.88, for MIQ scores see Figure 5A and B).

**Figure 5.**
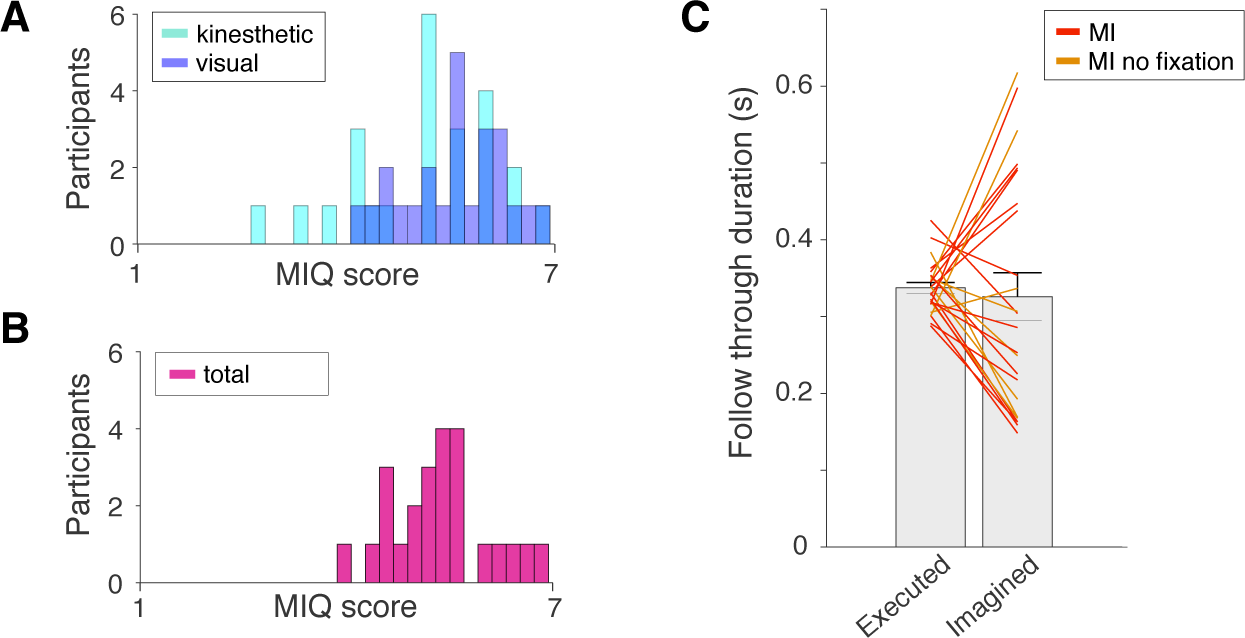
Measures of motor imagery. (A-B) Histograms of the MIQ scores of all motor imagery participants, divided into (A) measures of visual and kinesthetic imagery (B) and the total scores per participant. In each plot the range of possible scores (1-7) is bins into 30 units for visualisation. (C) Durations of executed and imagined follow-throughs for all participants in the motor imagery groups. The coloured lines indicate individual participants; bars show mean ± s.e. across participants

Although we did not directly measure muscle activity through EMG in our experiment, we could quantify net motor activity during motor imagery by overt movement. That is, subjects were required to be stationary at the central target during motor imagery and the handle of the vBOT was free to move so that any overt movement which would trigger a mistrial. During the exposure phase, the frequency of mistrials was not statistically different between the motor imagery group and the no motor imagery group for either breaks in fixation (6.0% and 5.3%, t(30) = 0.79, p = 0.44, for each group respectively), or central target overshoots (4.0% and 4.1%, t(30) = 0.26, p = 0.80). This suggests that neither type of mistrial was responsible for learning in the motor imagery group. We also compared various kinematic measures during pre-exposure to ensure that hand paths on the initial movement did not vary systematically with the secondary targets (see Supplementary Material).

## Discussion

Our results show that when subjects repeatedly reach in a force field whose direction can reverse from trial to trial, substantial interference occurs and there is no net learning. However, if subjects are asked to imagine making different follow-through movements for each field direction, they show substantial learning for identical movements. Critically, the only difference between these two conditions is motor imagery. This suggests that the act of imagining different future movements, even though subjects know they will not be performed, allows two motor memories to develop for the same physical state of the limb. In contrast to previous studies of mental practice, here imagery is not a mental rehearsal of the skill itself. That is, all subjects actually make the initial movement in the field, but the mental imagery is of a future movement (the follow through) which is separate from the skill itself. This suggests that knowing that you will imagine a subsequent movement changes the planning of the entire motor sequence so that the movement and motor imagery may be planned together prior to initiation. This supports a distinct role for mental imagery in the ability to learn novel skills.

Many studies have suggested that practicing a physical skill through motor imagery can result in improvement when subsequently performing the skill^22–25^. Traditional theories consider that such motor imagery acts as a simulator^26, 27^, whereby imagery can improve performance by using a forward model to predict the consequences of non-executed actions^23^. That is, a forward model allows a subject to try out different sequences of commands and compare the consequences, or to adapt a controller from the mentally simulated movement with the ensuing imagined error. The value of such a mechanism relies on the notion that, in general, forward models are easier to learn than controllers, as the desired output and the movement outcome can be compared to train a forward model during real action. In contrast the signal that is required to train a controller, that is the error in motor command, is not readily available^28^. Crucially, these studies of motor imagery consider the effects of mentally rehearsing the skill that is to be learned and typically compare learning under actual performance to either no practice or mental practice of the skill. Our study shows that subjects are able to learn two opposing skills, not by imagining the skills themselves, but by pairing each real execution of a perturbed physical movement with subsequent motor imagery that differs for each perturbation.

We have previously suggested that the key to representing multiple motor memories is to have each associated with a different neural state, rather than physical state of the body^1^. In this view, the interference seen in the no-imagery control group is due to repeatedly experiencing the same neural states for each reach to the central target. After each trial in an opposing field, the motor system will link these neural states to changes in the motor command, but over time these opposing adjustments cancel out, leading to no learning. Contexts which separate the neural states for the reach to the central target should allow learning by expanding the representation of the physical state to different neural states. This would then allow the same physical movement to be associated with different motor commands.

Taken together with previous results, our study suggests that distinct motor imagery and distinct motor planning can both lead to different neural states that can be used to form and retrieve different motor memories. However, this does not mean that motor planning and motor imagery are identical. Although both lead to equal amounts of learning, our data suggests only partial overlap between them as we observed incomplete transfer of learning from motor imagery to motor execution.

One way to create different neural states for the same physical state, is to change the context of each movement by making the movement to the central target part of a larger motor sequence. For example, the movement in the force field to the central target could have a different movement before^4^ or after it^1, 6^, enabling concurrent adaptation to force fields that would normally interfere. Alternatively, two recent studies have shown that adaptation to opposing force fields for spatially separate targets can be achieved for the same kinematic states, if visual feedback when reaching to each target is distinct. This maps the same kinematic states to different visual locations, producing different inferred states of the limb^5, 29^. In the dynamical systems perspective of motor cortex, planning sets an initial neural state, and execution arises from the subsequent evolution of the intrinsic neural dynamics^8, 30, 31^. Therefore, planning the same kinematic trajectory (movement to the central target) as part of a larger motor sequence will lead to a different initial neural state and a different subsequent neural trajectory. We have previously shown that planning different future movements, but aborting the plans before execution, allows learning of different force-fields over the same physical states. Here our results show that even when subjects know that they will not follow through, motor imagery of a follow through leads to the ability to learn opposing fields. This suggests that imagining different future movements may lead to distinct neural states from the start of the movement. Our hypothesis is consistent with recent electrophysiological work in non-human primates. Recently, Vyas and colleagues^15^ demonstrated that when monkeys used a brain-machine interface (BMI) to covertly rehearse cursor reaching movements, they adapted their cursor movements to visuomotor rotations, and moreover this adaptation transferred reliably but incompletely to overt arm reaching. Futhermore, the initial neural states for each centre-out BMI-controlled cursor movement closely resembled the initial neural states for the corresponding physical reaches. This consistency in neural dynamics between BMI-controlled and overt movement preparation is comparable to the learning and transfer observed here in humans instructed to imagine moving. Considering that similar motor cortical dynamical features are seen in humans and non-human primates^16^, this suggests that human motor imagery may evoke similar preparatory neural states to physical movement. In addition, human neuroimaging and electrocorticography studies have shown similar motor-related activity when imagining and executing movements^9–12, 32^ and similar effects on corticospinal excitability^33, 34^.

Our results demonstrate a complementary function for motor imagery. That is, in addition to its potential use as a simulator for possible actions, motor imagery can also engage distinct motor memories when preparing for the same physical movement. We show that mentally imaging a follow through movement can separate motor memories as well as actually performing or planning a follow through (Figure 3B). Moreover, such learning under mental imagery has significant transfer to full follow-through movements that are planned and then executed, as indicated by our measures of force adaptation on full follow-through channel trials. This suggests two features of motor imagery. First, that preparatory neural activity may be different when preparing to make the same movement when imagining different subsequent movements. Second, the generalization of this learning to physical action suggests that the neural states evoked when preparing for an imagined movement are similar to the states for the corresponding planned and executed movement.

The link between imagined and executed movement is supported by the similar chronometrics of the two. For example, imagined movements are known to have similar durations to executed movements^20, 21, 35, 36^ and show a speed-accuracy trade-off^21, 35, 37–39^. While the participants in our study demonstrated similar chronometrics for imagined and executed movements, the absolute difference between the time spent imagining and executing follow-throughs was uncorrelated with final adaptation. The amount of learning seen for each subject was also uncorrelated with performance on the MIQ-RS motor imagery questionnaire, used for assessing imagery ability^40^. However, a point to point movement is almost the simplest movement one can make, which, together with the relatively high mean scores (all above 3.9 out of 7, Figure 5A and B), may well have been simple to imagine.

It is still an open question as to what differences in neural state are necessary for the learning seen in our study. For example, it might require different activity in the same neural circuits or that different neural circuits are active. Recent studies of neural coding in motor cortex have emphasized its operation as a dynamical system in which planning involves setting the initial neural state and execution involves allowing the transitory dynamics to evolve from this state^8, 16, 41^. In this framework, our results would be accounted for by the same circuit but with different activity for each imagined movement. However, future work will be required to fully resolve how motor imagery leads to separate motor memories.

In summary, we show that simply imagining different future movements can enable the learning and expression of multiple motor skills executed over the same physical states. Our results suggest a new role for imagining in the representation of movement: to engage distinct motor memories for different future actions.

## Methods

We recruited 58 subjects (36 female; 25.0 ± 4.1 years, mean ± s.d.), with no known neurological disorders, who provided informed written consent and participated in the experiment. All participants were right handed according to the Edinburgh handedness inventory^42^ and were naive to the purpose of the experiment. The protocol was approved by the University of Cambridge Psychology Research Ethics Committee, and all experiments were performed in accordance with these guidelines and regulations.

Experiments were performed using a vBOT planar robotic manipulandum, with associated virtual reality system and air table^43^. The vBOT is a custom-built back-drivable planar robotic manipulandum exhibiting low mass at its handle. The position of the vBOT handle was calculated from optical encoders on the motors (sampled at 1 kHz). Endpoint forces at the handle of the robotic manipulandum were specified by sending commands to the torque motors. Participants grasped the handle of the vBOT with their right hand, with their forearm supported by an air sled which constrained movement to the horizontal plane. Visual feedback was provided using a computer monitor mounted above the vBOT and projected veridically to the subject via a mirror. This allowed us to display targets and a cursor representing the hand position (0.5 cm radius disk) overlaid into the plane of the movement.

In the fixation groups, eye movements were tracked using an SR Research Eyelink 1000 camera (sampled at 1kHz). The camera was positioned beneath a cold mirror. At the start of the experiment and after each rest break the eye tracker was calibrated over the visual work-space.

### Paradigm

Participants were divided into five groups. The number of subjects was chosen based on experience with similar motor learning experiments. Two subjects were excluded from analysis (see below) and replaced. Data from two of these groups (the follow-through and planning only groups) has been published previously (n=6 in each group) and is included here for comparison^1^. For these two groups we added two additional subjects to bring the number in each group to 8.

All participants made reaching movements in a horizontal plane from one of four starting locations to the central target (grey 1.25 cm radius disk), located approximately 30 cm below the eyes and 30 cm in front of the chest. The four starting locations were positioned 12 cm from the central target and arranged at 0° (closest to the chest), 90°, 180° and 270°. In addition to the start and central target, on each trial one of two secondary targets (yellow 1.25 cm radius disk) was displayed 10 cm from the central target and positioned at either +45° or −45° relative to the line connecting the starting and central target. The groups differed in whether they were required to continue the reach from the central target to the secondary target and what instructions they were given (see below).

During the movement to the central target the robot either generated no force (null field trials), a velocity-dependent force (exposure trials) or a simulated spring constraining the hand to a straight line path to the target (channel trials). Any movements from the central to the secondary target were made in a null field. On exposure trials the velocity-dependent curl force field was implemented as:

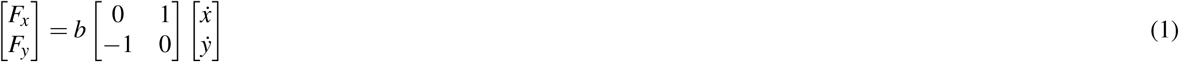

where 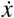 and 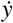 are the Cartesian components of the hand velocity and *b* is the field constant (± 15 N s m^−1^) whose sign determined the direction of the force field, that is clockwise (CW) positive and counter-clockwise (CCW) negative. The direction of the force-field applied during the movement to the central target was coupled to the position of the secondary target (e.g. CW for +45° and CCW for −45°). The association between secondary target position and curl field direction was fixed for each participant and counterbalanced across participants.

Channel trials were used to measure subject-generated forces, a measure of feed-forward adaptation^17, 18^. On a channel trial, the vBOT simulated a spring (spring constant of 6,000 N m^−1^ and damping coefficient of 50 N s m^−1^ both acting perpendicular to the wall) constraining the subject’s movement to a straight line to the central target.

#### Group 1: Follow through (n=8)

This experiment has been described previously^1^ and is included here for completeness. At the start of each trial, one of the starting locations was displayed and the hand was passively moved to it by the vBOT. The central target and one of the two possible secondary targets were then displayed (Figure 1A, Follow through). Subjects were required to remain within the start location for 300 ms, after which they were cued by a tone to start the movement and to move to the central target and then secondary target. Subjects had to remain within the central target for at least 50 ms before following through to the secondary target. Subjects were encouraged to make the entire movement between 400 and 800 ms. They received text feedback “correct speed”, “too slow” or “too fast” as appropriate. If subjects moved before the tone, took longer than 1.5 s to complete the movement, or took longer than 1 s to initiate movement after the tone, a mistrial was triggered and subjects were required to repeat the trial. At the end of each trial the vBOT passively moved the hand to the next starting location using a sinusoidal velocity profile.

A block consisted of 8 exposure trials and 2 channel trials, such that an exposure trial was experienced at each combination of the four starting positions and two possible secondary target positions (corresponding to the two different field directions). All channel trials were performed from the 0° starting position, one for each of the secondary target positions. The order of trials within a block was pseudorandom.

Before the experiment subjects were given 30 trials of familiarization in a null field. They then performed a pre-exposure phase of 5 blocks (40 null trials), an exposure phase of 150 blocks (1200 exposure trials), and finally a post-exposure phase of 3 blocks (24 null trials). Rest breaks (1.5 min) were provided approximately every 200 trials, with a longer rest break available in the middle of the experiment if required.

#### Group 2: Planning only (n=8)

The experiment has been described previously^1^. In the planning only group we isolated the effect of planning a follow-through without executing it. In contrast to the follow-through group, once the hand had moved 6 cm towards the central target, the secondary target was extinguished on all null and exposure trials (Figure 1A, Planning only). Participants were instructed that if the secondary target disappeared, they were not to execute the secondary movement but instead stop at the central target. We chose 6 cm so as to trade-off the length that we displayed the secondary target during the movement to the central target (as planning could take place during this movement) and the ability of participants to terminate the movement and not overshoot the central target by 3 cm, which would trigger a mistrial.

Critically, on all channel trials the secondary target did not disappear and subjects performed the full follow through. In order to encourage participants to plan the follow-through movement we included channel trials for all starting positions. Therefore, in this group we kept the total number of exposure trials the same as the follow-through group (1200 exposure trials), but doubled the number of channel trials, including them for each reach direction equally. Therefore a block was 12 trials including 4 channel trials. Across pairs of blocks we included two exposure trials and one channel trial for every combination of starting location and secondary target position, therefore we increased the pre-exposure phase to 6 blocks and the post-exposure phase to 4 blocks. Text feedback on movement duration was provided only on full follow through channel trials in order to match overall kinematics to the follow through group.

#### Group 3: Motor imagery (n=16)

In this group we examined the effect of imagining performing a follow through movement, with the knowledge that it would not be executed. In contrast to the planning only group, the central target colour (blue or grey) indicated whether participants had to execute a reaching movement and stop at the central target, or reach to the central and then the secondary target. When the central target was blue, they executed a movement only to the central target, but were asked to imagine making the follow through movement (Figure 1A, Motor imagery). When the central target was grey, participants planned and execute a full follow through movement. As in the planning only group, on follow through trials the movement to the central target was in a channel. On motor imagery trials participants were asked to press a button with their other hand to indicate when their imagined movement reached the secondary target. A single button was used for all secondary targets, so that the motor plan for this button press was not specific to the follow through direction. This enabled us to compare the duration of imagined and executed movements. Crucially for the imagery group, the hand was unconstrained when at the central target so any net motor activity would result in movement which we could observe.

This group were required to fixate on a small white cross located on the central target during each trial. This was to ensure that participants did not make eye movements to the secondary targets. Participants rested their forehead against a headrest and were required to fixate on the cross and maintain fixation within 3 cm for the duration of the trial. If subjects broke fixation or blinked, an error was triggered and the trial was repeated. Participants could move their eyes freely between trials.

Blocks were the same as for the planning only group. At the end of the exposure phase, a subgroup (half of participants) performed the post-exposure phase, as in the planning only group so that we could compare aftereffects. The other participants performed an additional 20 blocks (termed the probe phase) in which we assessed adaptation on motor imagery trials without follow through. In these probe blocks we kept the proportion of trials that were exposure trials the same as in previous blocks, but changed half of the full follow through channel trials to motor imagery channel trials (Figure 1B, middle). Therefore, these 20 blocks consisted of 160 exposure trials, 40 motor imagery channel trials and 40 full follow through channel trials. After the probe phase these participants performed the same post-exposure phase as the other participants.

During each rest break, participants were asked to evaluate their motor imagery in the previous set of trials (approximately 240 trials). They rated the ease with which they were able to imagine the movements (1-7, hard to easy scale, similar in style to the MIQ-RS^40^), and how frequently they imagined the movements (1-3 scale corresponding to ‘fewer than half the trials’, ‘most trials’ or ‘every trial’).

All participants also completed the MIQ-RS motor imagery questionnaire^40^ prior to the start of the experiment. This questionnaire has previously been evaluated for reliability and internal consistency of visual and kinesthetic measures of motor imagery^44^.

#### Group 4: No motor imagery (n=16)

This group was the same as the motor imagery group except that participants were not instructed to imagine making follow-through movements and did not press a button (Figure 1A, No motor imagery). To match the time spent at the central target with the motor imagery group, we made participants wait at the central target for the mean time it took them to execute the follow through movements on channel trials (the average of previous follow through trials). As in the motor imagery group, at the end of exposure phase half of participants performed the post-exposure phase to compare aftereffects. The other half of participants performed the same probe phase as in the motor imagery group, but without the use of motor imagery. Therefore participants performed no motor imagery channel trials (Figure 1B, right). After the probe phase these participants performed the same post-exposure phase as the other participants.

At the end of the experiment participants were asked if they had been imagining follow-through movements on trials where they had to stop at the central target. One participant responded that they had been, and was excluded from analysis and replaced by an additional subject.

One further participant was excluded from this group and replaced by an additional subject. Midway through the experiment, this replaced subject suddenly started producing a kinematic error in the direction opposite to the field and their adaptation measured on no motor imagery channel trials was greater than 6 standard deviations from the group mean.

#### Group 5: Motor imagery no fixation (n=8)

This group performed the same experiment as the motor imagery group but without constraints on their eye movements. This was to make the use of eye movements in this group comparable to the follow through and planning only groups. At the end of the exposure phase all participants immediately performed the post-exposure phase.

## Analysis

On channel trials we measured percent adaptation as the slope of the regression of the time course of the force that participants produced into the channel wall against the ideal force profile that would fully compensate for the field. To do this we extracted a 400 ms (or the maximum available) window of data centred on the time of peak velocity and calculated the force generated against the channel. We used the velocity along the channel to predict the force the vBOT would have applied on an exposure trial. We performed regression (with zero intercept) on these times series and expressed the slope as a percentage (slope of 1 = 100%), termed adaptation. We analyzed all channels trials for the follow through group, which were all performed from the 0° starting location. The other groups had channels trials for all starting locations and to match the number of channel trials analyzed we included all channels in the sagittal direction (0 and 180° starting locations). The inclusion of only 0° channel trials does not affect the statistical conclusions.

In addition, on null and exposure trials, we calculated the maximum perpendicular error (MPE) as the largest deviation of the hand from the straight line connecting the starting location to the central target. The sign of MPE on each trial was set such that a positive MPE indicated a kinematic error in the same direction as the force field (as would be expected in early learning).

To display hand paths (Figure 3), we extracted position data from when the hand left the start location until it entered the central target position. Each path was then linearly interpolated (*x* and *y* separately) so as to sample 1000 points equally spaced in time. For each participant, we considered trials at four different epochs of the experiment, which were the last block of the pre-exposure phase, the first and last blocks of the exposure phase, and the first block of the post-exposure phase. Therefore for each phase and participant we obtained a single hand path corresponding to each perturbation and start location. To generate a path for a group we calculated the average (and s.e.) of the participants’ paths and plot the mean with shading showing ± s.e.

To visualise the MIQ scores we plot the distributions of the average test scores, divided into questions on kinesthetic versus visual imagery (Figure 5A) and total average scores (Figure 5B). We binned the range of possible scores (1-7) into 30 units for visualisation.

For statistical analysis, our key measures were adaptation during exposure (as MPE can be affected by co-contraction) and MPE during post-exposure. We averaged adaptation and MPE for each subject within a block. To assess learning we used two measures. The first measure was the difference in adaptation between the average of the pre-exposure blocks and that of the final 6 blocks of exposure. The second measure assessed the aftereffects, calculated as the difference in MPE between the average of the pre-exposure blocks with the first two blocks of post-exposure. We use averaged performance per block in our evaluations of learning, as each start locations and direction of perturbation is tested once within a single block and we wish to assess net learning across these conditions. For both measures we performed a paired t-test across subjects for each group.

To compare learning between groups, we performed 3 planned comparisons of final adaptation on follow through trials (average of last 6 blocks of exposure) using unpaired t-tests. We perform multiple unpaired comparisons instead of a one-way ANOVA across groups as our study includes both controls that we expected to learn (follow through and planning only groups) and one which we expected to not learn (no motor imagery group). Therefore an ANOVA comparison of group means does not help to answer our question of whether or not motor imagery enables learning. Based on previous work we combined the follow through and planning only groups into a pooled follow through group. We compared the motor imagery group to the pooled follow through and no imagery groups. We also compared the motor imagery group to the motor imagery (no fixation) group. As these tests compare the learning that transferred to full follow-through trials, we performed an additional two comparisons to test learning independent of transfer. We compared the learning on motor imagery channels in the motor imagery group subjects who performed the probe phase (n=8, average of first 4 blocks of the probe phase), to the learning on no motor imagery channels in the no motor imagery group subjects who performed the probe phase (n=8, also the average of first 4 blocks of the probe phase), and to the final adaptation of the pooled follow through group. In addition, we performed unpaired t-tests comparing final adaptation between the subgroups of the motor imagery group and the no motor imagery group. We perform Shapiro-Wilk tests on these final adaptation data to test for normality.

We made several between-group comparisons using unpaired t-tests to compare different features of behaviour that could have affected learning. We performed between-group comparisons of the percentage of imagery mistrials due to breaks in fixation (excluding blinks or breaks made before the cue to move), hand overshoots of the central target, and of the time spent at the central target.

We report uncorrected p values.

### Assessments of motor imagery

For the groups who were asked to imagine the follow through movement, the duration of the imagined movement was taken as the time from reaching the central target until the button press. We used this to assess mental chronometry in each subject^20, 21^, which compares the average imagined movement duration to the average executed movement duration on the channel trials.

We regressed the percent adaptation of each subject in the final 6 blocks of the exposure phase against the absolute time difference between executed and imagined movements, and against three different self-reported motor imagery measures: MIQ score (1-7), average ease of imagery maintenance score (1-7) and average frequency of imagery score (1-3).

## Acknowledgements

We thank the Wellcome Trust, Royal Society (Noreen Murray Professorship in Neurobiology to D.M.W.), the Cambridge Commonwealth, European and International Trusts and the Rutherford Foundation Trust (H.R.S).

## Author contributions statement

H.R.S., J.N.I. and D.M.W. designed the study. H.R.S. and G.M.Z. performed the experiments. H.R.S. performed the analyses, and drafted the manuscript. H.R.S, J.N.I and D.M.W. edited the manuscript.

